# Community succession of the grapevine fungal microbiome in the annual growth cycle

**DOI:** 10.1101/2020.05.03.075457

**Authors:** Di Liu, Kate Howell

## Abstract

Microbial ecology is an integral component of wine production. From the vineyard to the winery, microbial activity influences grapevine health and productivity, conversion of sugar to ethanol during fermentation, wine aroma production, wine quality and distinctiveness. Fungi in the vineyard ecosystem are not well described. Here, we characterised the spatial and temporal dynamics of fungal communities associated with the grapevine (grapes, flowers, leaves, and roots) and soils over an annual growth cycle in two vineyards to investigate the influences of grape habitat, plant developmental stage (flowering, fruit set, veraison, and harvest), vineyards, and climatic conditions. Fungi were influenced by both the grapevine habitat and plant development stage. The core microbiome was prioritised over space and time, and the identified core members drove seasonal community succession. The development stage of veraison, where the grapes undergo a dramatic change in metabolism and start accumulating sugar, displayed a distinct shift in fungal communities. Co-occurrence networks showed strong correlations between the plant microbiome, the soil microbiome, and weather indices. Our study describes the complex ecological dynamics that occur in microbial assemblages over a growing season and highlight the importance of core community succession in vineyards. In addition to enriching our understanding of how plants and microbes interact, these findings may provide insights to craft wine regional distinctiveness and cope with global climate change.

## Introduction

The grapevine (*Vitis vinifera*) is naturally colonised by complex and diverse microorganisms, such as bacteria, filamentous fungi, and yeasts (Barata et al., 2012; Stefanini and Cavalieri, 2018), which substantially modulate vine health, growth, and productivity (Gilbert et al., 2014; Müller et al., 2016). Microbes originate from the surrounding environment (including soil, air, precipitation, animal vectors, native forests) and colonise both exterior (epiphytes) and interior (endophytes) (Goddard et al., 2010; Stefanini et al., 2012; Lam and Howell, 2015; Zarraonaindia et al., 2015; Morrison-Whittle and Goddard, 2018). This microbiota can be transferred to the grape must (or juice) and thus have a profound influence on wine composition, flavour, aroma, and quality (Ciani et al., 2010; Barata et al., 2012; Morrison-Whittle and Goddard, 2018). *Saccharomyces* yeasts (primarily *Saccharomyces cerevisiae*) and lactic acid bacteria (predominantly *Oenococcus oeni*) are primary drivers of wine fermentation and to form wine flavour and aroma (Swiegers et al., 2005). Beyond these species, our knowledge about the microbial ecology associated with grapevines and its role in the vineyard ecosystem is limited.

Recent studies posit the existence of geographically structured microbiota that contribute to the expression of regional distinctiveness of agricultural products, specifically, known as “*terroir*” in viticulture [reviewed by Liu et al. (2019)]. Significant biogeographic patterns are generally observed at scales of hundreds to thousands of kilometres (Gayevskiy and Goddard, 2012; Bokulich et al., 2014; Taylor et al., 2014; Knight and Goddard, 2015), and influenced by multiple factors, such as climate and topography, soil properties, and even local anthropogenic practices (Bokulich et al., 2014; Pinto et al., 2014; Burns et al., 2015; Zarraonaindia et al., 2015; Jara et al., 2016; Portillo et al., 2016; Morrison-Whittle and Goddard, 2017). Microbial spatial pattern analyses have mainly been conducted at large scales and focused on studying grapes and must, the communities present at the start of fermentation, and the soil, but the nature and the importance of microbial ecology within vineyards towards defining *terroir*, is largely unknown. Our recent study demonstrates that vineyard ecosystems are structured by microbial communities and impact wine aroma and regional characteristics at the scale of 400 km, in particular by grapevine-associated fungi (Liu et al., under review).

Metagenomics tools have been actively used to map the plant microbiome, in particular the phyllosphere and the rhizosphere (Turner et al., 2013; Müller et al., 2016). In viticulture, many studies have been dedicated to characterising grapevine-associated bacteria and their relevant importance on plant growth and winemaking processes (Zarraonaindia et al., 2015; Marasco et al., 2018). Zarraonaindia et al. (2015) investigated the spatial and temporal dynamics of the bacterial microbiota associated with grapevine organs of Merlot cultivars in New York state, and revealed significant differences among grapevine habitats and the soil, although the latter serving as a source reservoir for the former. The endosymbiotic mycorrhizae are diverse and structured in rhizocompartments and the vascular system (Deyett and Rolshausen, 2020). Fungal communities across multiple compartments have not been systematically investigated. As described in other plant systems, microbiome composition is not temporally stable but change in response to plant development (Mougel et al., 2006; Jumpponen and Jones, 2010; Chaparro et al., 2014; Copeland et al., 2015; Grady et al., 2019) and can fundamentally affect the health and productivity of plants. Core microbiomes are commonly defined based on abundance-occupancy distributions describing microbes which are highly abundant and ubiquitous occurring the habitat over time and are hypothesised to reflect underlying functional relationships with the host and therefore important for managing plant microbiome towards desired outcomes (Lundberg et al., 2012; Shade and Stopnisek, 2019). In grapevines, however, this remains to be shown.

In the present study, we sampled microbial communities associated with Pinot Noir grapevines from two vineyards within the same grape growing region to provide insights into fungal ecology in the vineyard. Using internal transcribed spacer (ITS) amplicon sequencing to characterise fungal communities, we disentangled the influences of geographic location (between two vineyards), grapevine habitat (soil, roots, leaves, flowers and grapes), grapevine developmental stage (flowering, fruit set, veraison, harvest), and climatic conditions on the fungal diversity, composition, and structure from root zone soil, roots, leaves, and grapes or flowers. Specifically, we aimed to prioritise the core microbiome over space and time, and further confirm the role of this core community to structure seasonal community succession based on Random Forest supervised learning models. The influence of the changing environment was elucidated with network analysis. We show that fungal communities differ according to grapevine habitats and developmental stages, with little geographic influences at this scale (5 km). The core microbiome correlates closely with solar radiation and water status during the annual growth cycle and drives seasonal community succession. Combining our study with studies in other grape growing areas will expand our knowledge of the core microbiome associated with grapevine across different soil types and environmental conditions to allow optimisation of the growth of this important economic crop.

## Results

### Fungal communities vary by grapevine habitat within vineyards

A total of 4,786,543 ITS high-quality sequences were generated from all samples (n = 160), which were clustered into 2,286 fungal OTUs with a threshold of 97% pairwise identity. Among all the OTUs identified in the present study, 62.90% were detected in more than one grapevine habitat (root zone soil, root, leaf, flower and grape), of which 54 OTUs were shared across sample types (Fig. 1A). The soil contained the most OTUs and of these, 37.67% were also found associated with the plant, and 32.75% of soil OTUs were shared with roots which were higher than the diversity found in other plant organs. Each plant organ shared the majority of fungal taxa with other organs and the soil. Some OTUs were unique, for example, 28.11% of OTUs were observed exclusive to grapes (Fig. 1A). *Ascomycota* was the most abundant phylum in the vineyard habitats comprising 77.69% of all sequences, followed by *Basidiomycota* and *Mortierellomycota* in the plant, and *Mortierellomycota* and *Basidiomycota* in the soil, respectively (data not shown). The major fungal genera were *Aureobasidium, Cladosporium, Epicoccum, Mortierella, Cryptococcus, Debaryomyces, Saccharomyces, Mycosphaerella, Lophiostoma, Alternaria*, and *Penicillium*, with relative abundance more than 1.0% across all samples (Fig. 1B). Some major taxa associated with plants, such as *Aureobasidium, Debaryomyces, Saccharomyces* and *Mycosphaerella*, were present in soil samples at very low abundances (< 0.05%). The genus *Mortierella* had the highest relative abundance (26.00%) in the soil, and showed significantly lower abundances (ANOVA, F = 4.325, *p* < 0.01) in the plant (Fig. 1B).

**Fig. 1.**
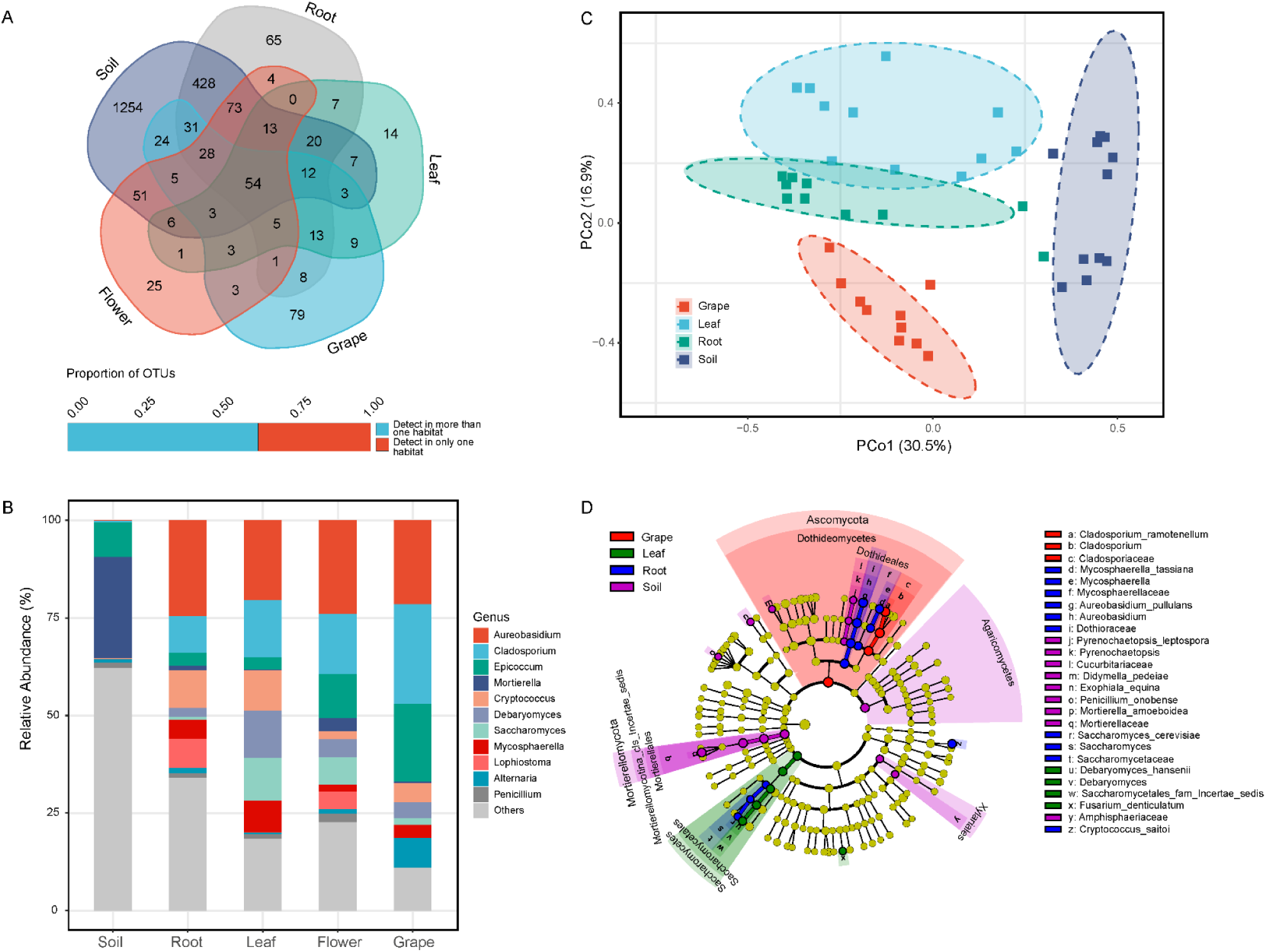
Fungal communities demonstrate different structures and compositions depending on habitats in the vineyard. (A) Fungal OTUs shared among soil and grapevine habitats (roots, leaves, flowers, grapes) in the vineyard; (B) Microbial community composition characterised to the genus level (relative abundance > 1.0% shown); (C) Principal coordinate analysis (PCoA) among plant organs and soil samples at harvest stage based on Bray-Curtis distances; (4) Linear discriminant analysis (LDA) effect size (LEfSe) taxonomic cladogram identifying significantly discriminant taxa associated with habitats at harvest.

Grapevine-associated fungi varied with vine habitat (root zone soil, root, leaf, flower and grape) and exerted significant differences in both community richness and diversity (*α*-diversity, Shannon index; ANOVA, F = 5.206, *p* < 0.001) and taxonomic dissimilarity (*β*-diversity, Bray–Curtis distance; ADONIS, R^2^ = 0.271, *p* < 0.001) independent of phenological development stage (Table 1). This trend was more distinct with an improved coefficient of determination (R^2^) within a certain stage. Geographic location only weakly impacted microbial diversity and composition, except for fungal diversity associated with roots, which was significantly impacted by vineyard (ANOVA, F = 5.878, *p* < 0.01) (Table 1). Principal coordinate analysis (PCoA) showed the habitat-specific patterns (95% confidence interval) of fungal communities, with 47.4% of total variance explained by the first two principal coordinate (PC) axes using the harvest stage as an example (Fig. 1C). Linear discriminant analysis (LDA) effect size (LEfSe) analysis further confirmed that these patterns related to significant associations (Kruskal–Wallis sum-rank test, *α* < 0.05) between fungal taxa and sample types (Fig. 1D). At harvest, *Mortierellomycota* (notably *Mortierella amoeboidea*), *Agaricomycetes, Xylariales* (including *Amphisphaeriaceae*), *Cucurbitariaceae* (including *Pyrenochaetopsis*), *Didymella pedeiae, Exophiala equina* and *Penicillium onobense* were observed with higher abundances in the soil. For grapevine habitats, *Saccharomycetaceae* (notably fermentative yeast *S. cerevisiae*), *Cryptococcus saitoi* (*Basidiomycota* yeasts), *Dothioraceae* (including *Aureobasidium*) and *Mycosphaerellaceae* (including *Mycosphaerella*) were significantly abundant in roots, with *Saccharomycetes* (including *Debaryomyces*) and *Fusarium denticulatum* in leaves, and *Ascomycota* (notably *Cladosporium*) in grapes.

**Table 1.** Experimental factors predicting α- and β-diversity of fungal communities in the vineyard

| Factor | Sample Group | α-diversity (Shannon)<sup>1</sup> | | β-diversity (Bray–Curtis)<sup>2</sup> | |
| --- | --- | --- | --- | --- | --- |
|  |  | F | *p* | R<sup>2</sup> | *p* |
| Habitat | All stages | 5.206 | **1.05E-02** | 0.271 | **0.001** |
|  | Flowering | 18.109 | **4.00E-11** | 0.366 | **0.001** |
|  | Fruit set | 38.109 | **4.00E-11** | 0.385 | **0.001** |
|  | Veraison | 5.668 | **2.93E-03** | 0.466 | **0.001** |
|  | Harvest | 22.848 | **2.27E-08** | 0.350 | **0.001** |
| Stage | Soil | 3.858 | **0.018** | 0.229 | **0.001** |
|  | Root | 1.390 | **0.026** | 0.155 | **0.008** |
|  | Leaf | 3.396 | **0.001** | 0.120 | **0.014** |
|  | Grape | 6.392 | **2.23E-02** | 0.426 | **0.001** |
| Vineyard | Soil | 3.969 | 0.055 | 0.034 | 0.203 |
|  | Root | 5.878 | **0.008** | 0.052 | 0.073 |
|  | Leaf | 0.018 | 0.894 | 0.017 | 0.806 |
|  | Flower | 0.988 | 0.353 | 0.513 | 0.757 |
|  | Grape | 0.788 | 0.382 | 0.025 | 0.551 |

<sup>1</sup> One-way ANOVA <sup>2</sup> Permutational multivariate analysis of variance using distance matrices Bold values indicate statistically significant results, *p* < 0.05

### Temporal fungal community dynamics at grapevine phenological stages

Significant variances in the fungal richness and diversity were recorded at each grapevine habitat over time (Table 1, Fig. 2A). Soil fungi showed higher diversity than that of grapevine organs (except flowers) throughout the growing season, with a rapid increase before fruit set, and then a decrease by veraison, and finally an increase at harvest. The trend was also observed in root fungal richness and diversity. After fruit set, the richness and diversity of grape fungi significantly increased over the development and ripening process, reaching a maximum at harvest. For leaves, the fungal communities tended to be less complex and diverse as grapevine development progressed, although an increased Shannon index was observed at veraison. Interestingly, veraison was found to be a key stage of diversity changes in grapevine-associated fungi microbiota (Fig. 2A). There were significant differences in fungal community composition over time regardless of grapevine habitat (Table 1). We observed the most distinct pattern in grapes with the highest R^2^ coefficients (ADONIS, R^2^ = 0.426, *p* < 0.001), followed by the soil (ADONIS, R^2^ = 0.229, *p* < 0.001). As an example, PCoA showed a strong separation of grape samples, especially samples at harvest from earlier stages on PC1, which explained 54.9% of the variance (Fig. 2B).

**Fig. 2.**
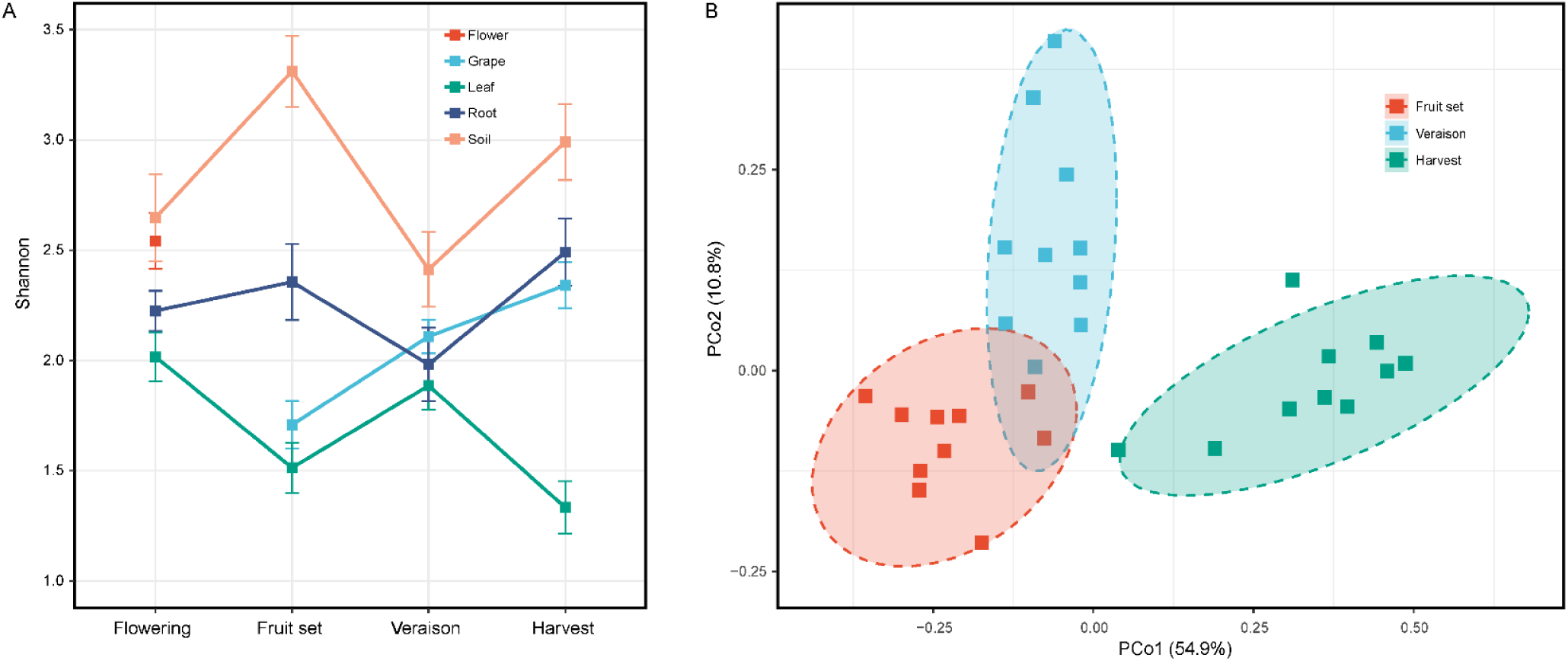
Grape developmental stage influences microbial diversity in the vineyard. (A) *α*-diversity (Shannon index) changes during the growing season (bars indicate SE); (B) Bray-Curtis distance PCoA of grape fungal communities according to the developmental stage. Shown are grapes sampled from fruit set, veraison, and harvest stage. Flowering was excluded as flowers were sampled and counted as another grapevine habitat.

### The core microbiome fluctuates according to plant development

Here, we prioritised the core microbiome over space and time to further investigate fungal community succession in the vineyard. We first identified the most dominant OTUs based on the abundance-occupancy distributions (see Experimental procedures for details) to include in the core (see Table S1 for a complete list) for the grapevine and the soil, respectively. About 1.51% of OTUs (15 OTUs) constituted a core that accounted for 74.70 ± 2.70% of the sequences across all plant samples (Fig. 3A; Table S1A). On average, the core community was highly abundant in each grapevine habitat, with a maximum recorded in grapes (90.55 ± 2.27% in relative abundance) (Fig. 3B; Table S1A). A wide variety of fungi was observed within those core members, and included fermentative yeasts (*Saccharomyces, Debaryomyces*), yeast-like fungi (*Aureobasidium, Cryptococcus, Vishniacozyma*), filamentous fungi (*Cladosporium, Alternaria, Penicillium, Fusarium*), and other genera (*Mycosphaerella, Didymella, Ramularia, Epicoccum*) (Table S1A). Within grapevine habitats, some species outside the grapevine core dominated and developed over time and also defined the core specific to this organ (Table S1A1 – A4). Specifically, three species from *Mortierella* genus and *Microascaceae* family were as part of core community of flowers (Table S1A1), with three from *Filobasidium* and *Alternaria* genera for grapes (Table S1A2), and three from *Hypocreales* order, *Thermoascaceae* family and *Lophiostoma* genus for roots (Table S1A4). For leaves, only eight OTUs within the grapevine core dominated the community, while other core OTUs showed lower occupancy and/or abundance during the growth cycle (Table S1A3). For soil, 1.74% of OTUs (35 OTUs) were the core members accounting for 63.40 ± 3.54 % of the reads across all samples, which presented a high abundance at each stage (Table S1B). The core taxa belonged to *Ascomycota, Mortierellomycota* (*Mortierella*), *Basidiomycota* (for example *Clavaria* and *Auricularia*) and *Chytridiomycota* (*Rhizophlyctis*), including some filamentous fungi like *Podospora, Humicola, Ilyonectria* and *Sarocladium*, and yeasts including *Exophiala* and basidiomycetous *Solicoccozyma*.

**Fig. 3.**
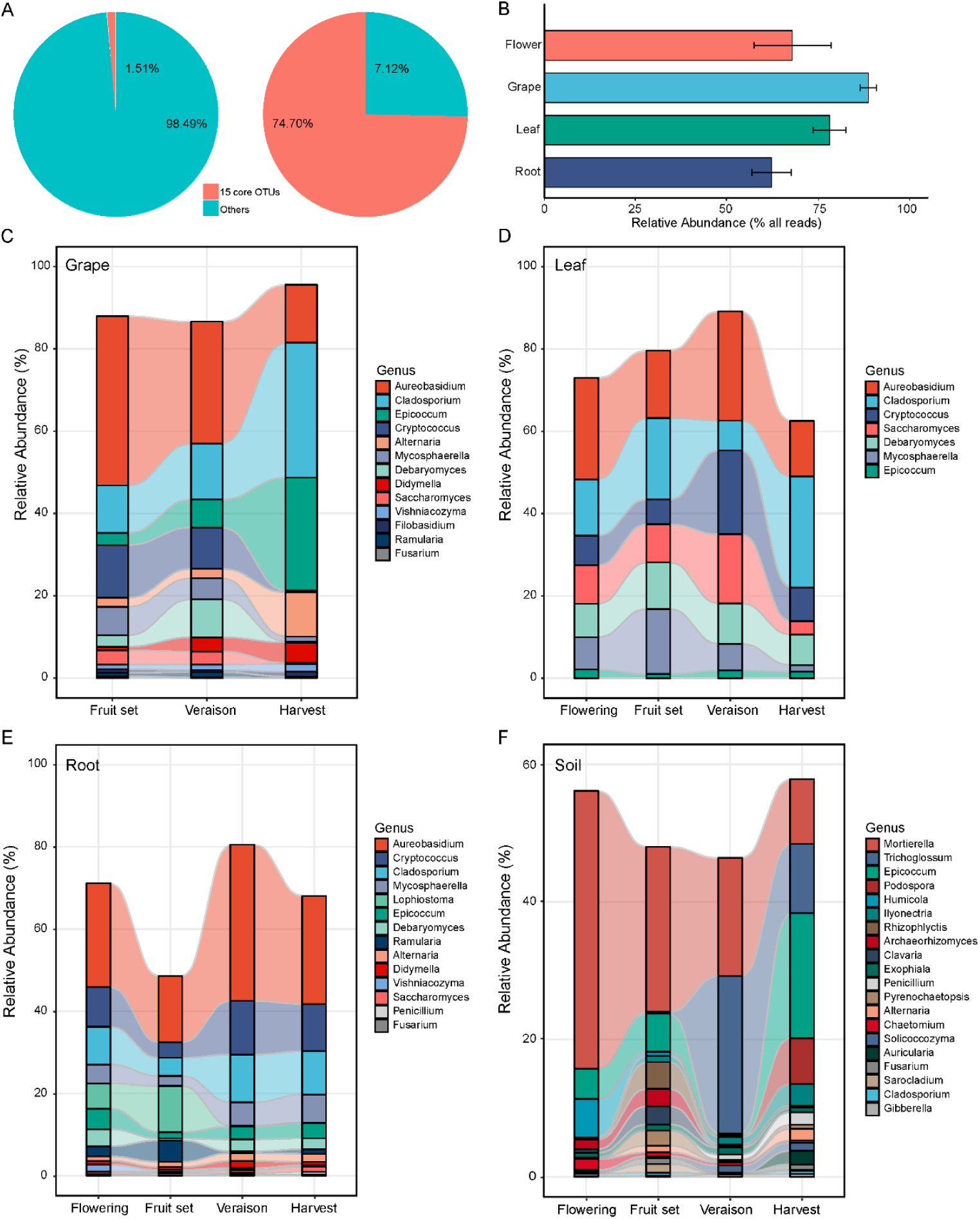
The core microbiome in grapevine habitats and the soil over the growing season. (A) Percentage (left) and relative abundance (right) of OTUs representing the core versus the remaining microbiome associated with the grapevine (flowers, grapes, leaves, roots); (B) Relative abundance (mean ± SE) of dominant OTUs across plant habitats; (C – F) Relative abundance changes of dominant fungal genera of grapes (C), leaves (D), roots (E), and soil (F) from the beginning to the end of growing season.

Tracking core OTUs within individual habitats revealed distinct temporal dynamics and succession associated with plant development (Fig. 3C – F). All the core genera in the alluvial diagrams presented significantly different relative abundances during the annual growth cycle (ANOVA, *p* < 0.05). Overall, veraison appeared to be a key stage where the core community was distinct from the others across habitats (Fig. 3C – F). Some taxa changed in a similar manner in plant organs. For instance, the most dominant genera, *Cladosporium*, accumulated in relative abundance, between post-veraison grapes and leaves, while, *Aureobasidium* and *Cryptococcus* declined in those post-veraison organs (Fig. 3C – E). Fermentative yeasts *S. cerevisiae* and *Debaryomyces hansenii* persisted in the grapevine throughout the growth cycle but decreased in relative abundances of post-veraison grapevine habitats. However, *S. cerevisiae* increased in the root microbiome post-veraison. Some species like *Epicoccum* sp. and *Alternaria* sp. showed changes in developing grapes and became much more abundant at harvest, but these genera only showed moderate changes in leaves and roots (Fig. 3C – E; Table S1A2 – A4). It is noteworthy that a few core members only appeared or appeared in high occupancy in specific developmental stages. The best example is OTUs assigned to *Alternaria infectoria* and *Alternaria rosae* where taxa only were only detected in grape samples at veraison and accumulated afterwards (Table S1A2). For the soil habitat, the most dominant genus *Mortierella* decreased in the relative abundance during the plant development (especially after veraison), accompanied by an increase in *ascomycetous* genera (Fig. 3F). Within ascomycetous genera, those taxa shared between soil and plants accumulated as the growth cycle processed, including *Epicoccum, Alternaria, Fusarium, Cladosporium* and *Penicillium*, with notably great changes at veraison, as well as soil yeasts *E. equina* and *Solicoccozyma phenolica*, and some soil-specific fungi, such as *Trichoglossum, Podospora* and *Humicola* (Fig. 3F; Table S1B).

### Core microbiome drives seasonal community succession

To further confirm the robustness of these observations across the growth cycle, Random Forest supervised learning models (Breiman, 2001) were employed to classify samples and identify which taxa explain the strongest variations during grape vine developmental stages. The fungal microbiota had high discriminative power to distinguish samples coming from various stages, especially for grapes and soils (class errors, 0.100 and 0.125 on average, respectively; Table S2). Compared to all other stages, veraison had the lowest predictive error (class error = 0.150 on average), for example, all soil samples at veraison were correctly identified (class error = 0.000), suggesting that veraison was the most distinct stage of the development.

All the core OTUs were important features of these models for grapevine habitats (Fig. 4). Filamentous fungi (for example *Cladosporium ramotenellum* and *Fusarium denticulatum*), fermentative yeasts *S. cerevisiae* and *D. hansenii*, and some yeast-like fungi (*Aureobasidium pullulans* and *Cryptococcus saitoi*) within the core (Table S1) were top features in the classification models for vine habitats. Several non-core taxa were also important for the models (see Table S3 for a complete list). For instance, yeast *Hanseniaspora uvarum* (OTU_33) presented with low occupancy (≤ 50%) across organs with higher abundance at veraison and harvest as were yeasts *Torulaspora delbrueckii* (OTU_121) for both grapes and leaves, and *Rhodotorula babjevae* (OTU_40) for grapes. In addition, some soil core species were important for plant models, including *Mortierella* spp., *Sarocladium bactrocephalum, E. equina*, which were only presented at specific stages (Table S2B). A portion of soil core taxa explained the highest variations of some phenological stages, for instance, *Epicoccum nigrum* (OTU_15) and *Pyrenochaetopsis leptospora* (OTU_62) were the most important predictors (Fig.4; Table S1). Non-core taxa were mainly detected at early stages (flowering and fruit set; e.g., *Archaeorhizomyces* sp., *Solicoccozyma terricola*) or later stages (e.g., *Penicillium multicolor, Aspergillus ardalensis*), and contributed to the models. When we used the core subset to build the random forest model, a higher classification accuracy (class error = 0.075 on average) was achieved, with improved discrimination for fruit set and harvest (0.200 and 0.000, respectively). Taken together, these results confirmed that plant development stages can be distinguished based on the associated core microbiome and defined predictive features for dynamic fungal ecology in the vineyard.

**Fig. 4.**
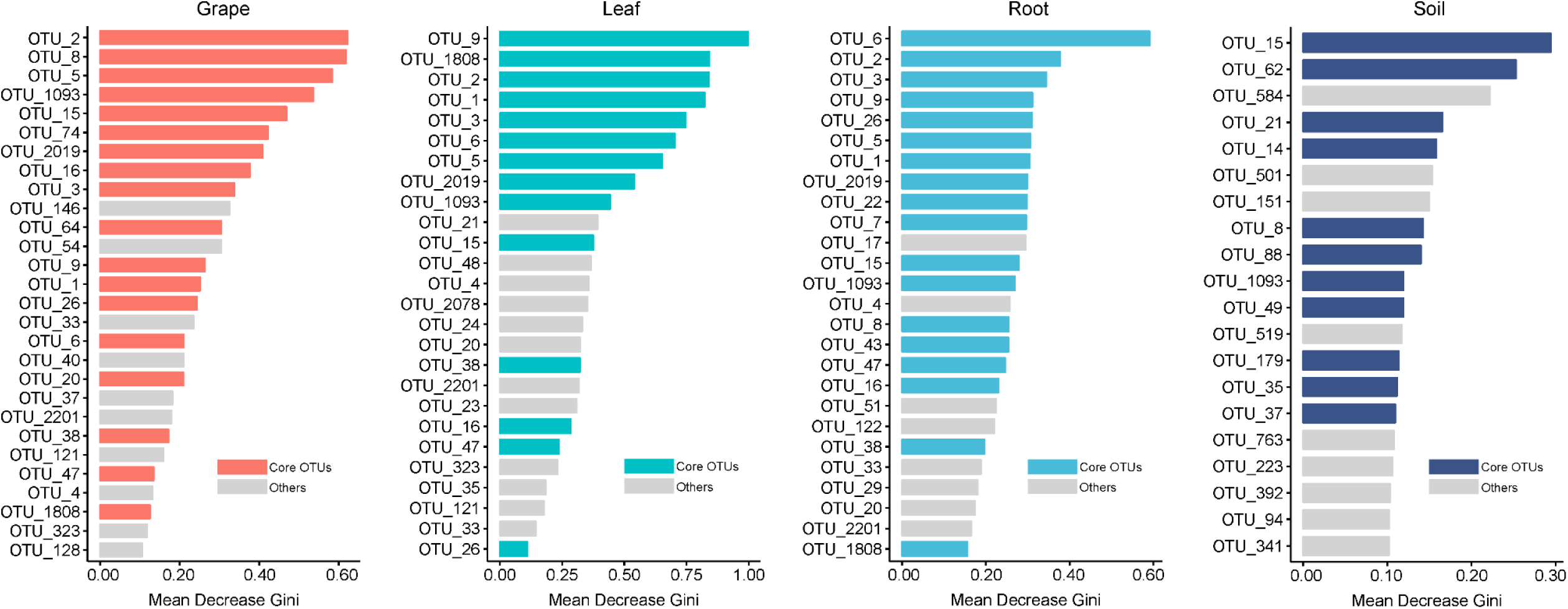
The core microbiome (as OTUs) can distinguish developmental stages for plant habitats and soil. Shown are the important features (values > 0.10) based on Mean Decrease Gini (MDG) of random forest models. Core members were coloured in pink (grape), green (leaf), blue (root), and navy (soil). Identified OTU species are given in supplementary table S1.

### Co-occurrence patterns between the core microbiome and vineyard weather

During the annual growth cycle, all weather parameters displayed significant differences between the phenological stages (Supplementary Table S4). Stage-dependent patterns in grapevine-associated microbiota also suggest that longitudinal environmental conditions are likely responsible for structuring the microbial communities. Network analysis was conducted to explore the co-occurrence patterns between plant core microbiome and environmental matrices including abiotic (weather) and biotic factor (soil microbiota), based on the strong correlation coefficients (Spearman correlation coefficient, ρ ≥ 0.8; p < 0.01) (Fig. 5). Overall, the plant microbiota presented high connectivity with soil microbiota and weather conditions. The more densely connected module was observed in grape microbiota (average degree = 3.610; Fig. 5A) than leaf (average degree = 2.381; Fig. 5B) and root (average degree = 2.316; Fig. 5C). In addition, grape microbiota was significantly correlated with each weather parameter. Across habitats, the most closely connected fungi were *A. pullulans* (OTU_1), *C. ramotenellum* (OTU_2), *C. saitoi* (OTU_3), and *F. denticulatum* (OTU_2019). Correspondingly, high-connectivity nodes with these OTUs were soil species *Penicillium* sp., *Auricularia* sp., *Humicola* sp., *Mortierella* sp., and *Ilyonectria* sp., and changing solar radiation (Env1) and vine water status related parameters (Env6, evaporation; Env8, relative soil moisture). Likewise, soil taxa were also strongly correlated with weather parameters (in particular Env1, 6, 8) (Supplementary Fig. S1). In addition, habitat-specific correlations were also found. For example, grape-specific core taxa *A. infectoria* (OTU_74) and root-specific core taxa *Lophiostoma* sp. (OTU_22) were densely connected nodes for them, respectively. The below ground root microbiota were significantly correlated with relative soil moisture.

**Fig. 5.**
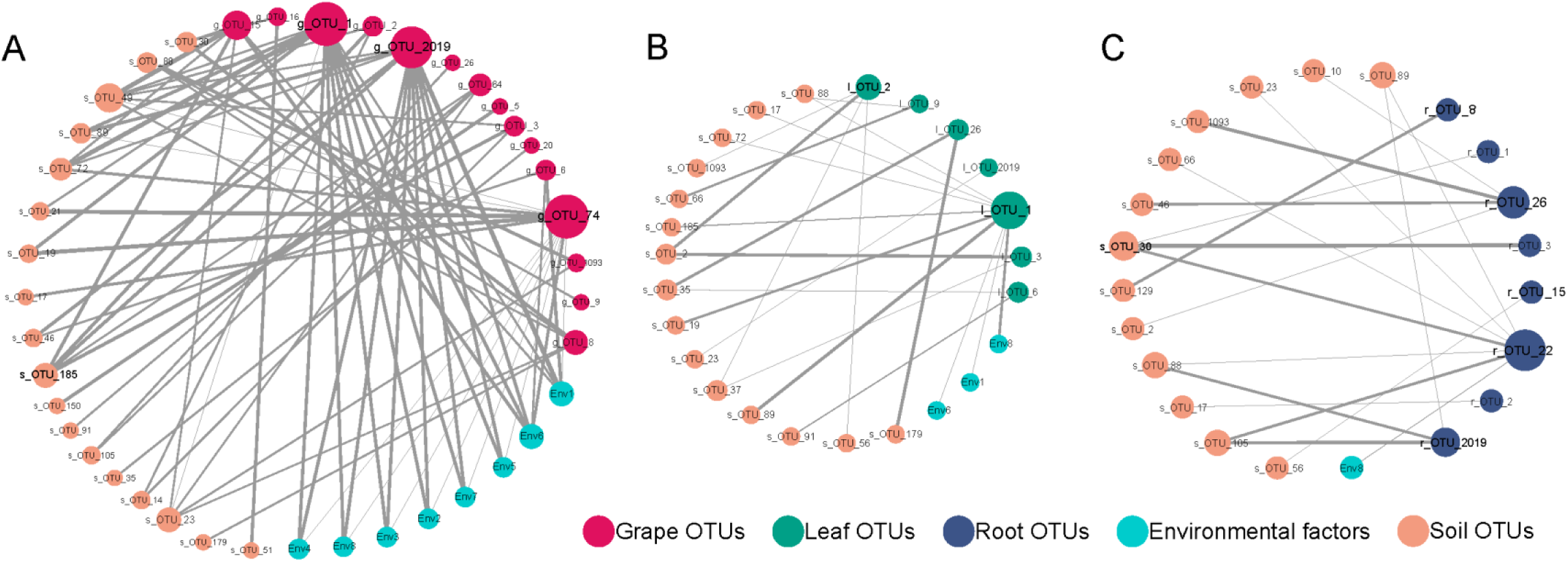
Statistically significant and strong co-occurrence relationships among grapevine-associated and soil dominant fungi and weather conditions during annual growth cycle. Network plots for grape (A), leaf (B), and root (C) core microbiota with weather matrices and soil core microbiota. Circle nodes represent core OTUs and environmental variables, with different colours. Direct connections between nodes indicate strong correlations (Spearman correlation coefficient, ρ ≥ 0.8; p < 0.01). The size of nodes is proportioned to the interconnected degree. The width of edges is proportioned to the correlation coefficients. Abbreviations: Env1, solar radiation; Env2, mean high temperature; Env3, mean low temperature; Env4, mean temperature; Env5, precipitation; Env6, evaporation; Env7, transpiration; Env8, relative soil moisture.

## Discussion

Microbiomes are dynamic over space and time. Despite the overwhelming diversity of microbial communities, relatively few microbial taxa dominate the ecosystems locally or globally (Delgado-Baquerizo et al., 2018; Egidi et al., 2019; Grady et al., 2019; Shade and Stopnisek, 2019). Describing the core microbiome is essential for unravelling the stable, consistent components across complex microbial assemblages. The definition of a core microbiome provides the scope to define a reproducible and conservative perspective for the dynamic microbiome, and allows for systematic discovery of persistent taxa, especially for microbiota associated with perennial plants (Shade and Stopnisek, 2019). Rare taxa that conditionally (here, specific to the habitat and the development stage) contribute to changes in microbial diversity would provide a different and arguably less ecologically relevant community for perennial crops (Shade et al., 2014; Shade and Gilbert, 2015). Here, our study narrows down the immense number of measurable fungal taxa to the core microbiome to advance understanding of grapevine-associated fungal microbiota. This is an important step to further investigate contributions of fungi to functioning of the vineyard ecosystem and opens the potential to use specific fungi to deliberately manipulate the microbiome to affect grapevine health, growth, and grape and wine production.

### Community succession of the grapevine-associated microbiota over space and time

Our data show that fungal communities are not stochastically assembled, but have a distinct community with habitat patterns according to the development stage of grapevines (Fig. 1). Fungal communities varying by sites have previously been shown in vineyard systems (Coller et al., 2019; Knight et al., 2019). In our study, grapevine habitats and plant development within the sampled vineyards outweighed biogeographic trends, which were only significant for root fungal richness, suggesting that fungal assemblages have extensive local heterogeneity associated with their host plant (Table 1). Grapevines maintain distinct fungal microbial communities in a habitat-specific manner, but also share many taxa. Fungal diversity is higher below the ground (root zone soil, roots) than the aboveground habitats (flowers, leaves, and grapes), and this is in line with other studies on grapevines (Zarraonaindia et al., 2015; Morrison-Whittle and Goddard, 2017; Deyett and Rolshausen, 2020). Soil has been suggested as the source and reservoir of plant microbiota (Zarraonaindia et al., 2015; Morrison-Whittle and Goddard, 2018; Grady et al., 2019) yet many core taxa associated with grapevine organs (for example, *Cladosporium, Aureobasidium*, and *Saccharomyces*) are in low abundances in our soil samples (Fig. 1B, 1D). This differentiation indicates that there are ecological filters operated by the plant to favour some species while excluding others (Chesson, 2000; Leibold and McPeek, 2006). The core taxa found here persist across habitats in differing abundance with some species only existing to a specific plant organ. We observed a decrease in fungal diversity between the root zone soil and the roots but an enrichment of saprophytic fungi (*Mycosphaerella* and *Lophiostoma*, core taxa in roots; Fig. 3E) and dilution of potentially pathogenic fungi (*Ilyonectria*, core taxa in soil; Fig. 3F). These fungi have been previously reported colonising the rhizocompartments of grapevines (del Pilar Martínez-Diz et al., 2019; Deyett and Rolshausen, 2020). Arbuscular mycorrhizal fungi including *Glomeromycota*: *Rhizophagus, Funneliformis*, and *Acaulospora*, which positively affect grapevine growth in a mutualistic manner (White, 2015), were recovered with low prevalence and abundance in root zone soil and root samples in our study (data not shown). This could be due in part to the choice of barcoding region used here, rather than the large subunit rRNA, which is the preferred biomarker for these fungi (Lekberg et al., 2012). However, geographic location may also influence distribution (Kivlin et al., 2011; Hazard et al., 2013), as found in vineyards by Coller et al. (2019) who retrieved *Glomeromycota* as a core phylum in Italy, using the same ITS1F/2 primers as in our study. Interestingly, *Mortierella*, a zygomycete genus and an important component of phosphorus cycling in the rhizosphere (Osorio and Habte, 2001), were observed as core members associated with both soil and flowers (Fig. 1B; Supplementary Table S1). The presence of this genus in flowers is unexpected, and may point to a role of flowers in forming and establishing the grapevine microbiome and their functionality.

The grapevine-associated microbiota are also affected by the plant development stage. The leaf fungal diversity decreased during the growth cycle (Fig. 2A), as previously described for grapevine and other plant systems (Pinto et al., 2014; Copeland et al., 2015). This may be partly attributed to experiencing greater climatic stress (UV radiation, temperature, and humidity) in the phyllosphere and thus only eight core OTUs persist over time due to environmental fluctuations. Microbial richness and diversity increased in the grapes from fruit set to harvest stage (Fig. 2A), and may be explained by sugar accumulation and exudation as ripening proceeds. Accumulation of sugars favours fungal colonisation (Martins et al., 2014). Similar trends between soil and root fungal diversity were as expected, suggesting that root microbiomes are partly derived from the rhizosphere (Compant et al., 2011; Lundberg et al., 2012; Deyett and Rolshausen, 2020), and in turn, that root morphology and root exudation can influence the composition of the rhizosphere microbiome (Berg and Smalla, 2009; Chaparro et al., 2014).

Our study shows that the core microbiome of grapevines exists over space and time in the vineyard ecosystem. The increase in the relative abundance of soil core taxa that is shared with plants over the growing cycle lends further support to the influence from the roots on plant development (Fig. 3F). *Aureobasidium* are the most widespread fungi in the vineyard, and persist throughout the growing season and decline at later stages across plant organs (Fig. 3), supporting previous studies in vineyard ecosystems (Renouf et al., 2005; Martins et al., 2014; Pinto et al., 2014). Using high-throughput sequencing, our data reveal the persistence of *S. cerevisiae* across habitats and developmental stages as core taxa, which has previously been detected in mature grapes based on enrichment culture-dependent methods (Fleet et al., 2002; Mannazzu et al., 2002; Renouf et al., 2005) and at low frequencies in leaves (Pinto et al., 2014). We did not observe an expected increase in the relative abundance of *S. cerevisiae* after veraison in grape and leaf samples (Fig. 3C, 3D), but there could be increased competition with other species, for example *Cladosporium* as sugar becomes more available. Meanwhile, some non-core OTUs displayed temporal succession, which were screened by a supervised learning approach (Fig. 4). These predictors associated with grapevine displayed seasonal volatility, for instance, *Filobasidium stepposum, E. equina*, and *Clonostachys rosea* were not detected at veraison yet reappeared at harvest in roots, as did *M. amoeboidea* for leaves, and *S. bactrocephalum* for grapes (Fig.4; Supplementary Table S3). Cyclic patterns have been reported in the human gut microbiome and indicate that some dynamic lineages of microbes decreased in prevalence and abundance in modernised populations (Smits et al., 2017). In this study, those non-core OTUs were not considered as the main driver of seasonal community succession due to lack of persistence and clear biological functions, but their dynamics might indicate that there is some adaptation process for the plant or plant-microbe interactions. Further research will determine if this volatility has an ecological or functional role in the vineyard.

Veraison is the stage where the fungi microbiome of the grapevine changes very distinctly. In grapevine phenology, veraison marks the beginning of ripening and is a crucial time point of grapevine metabolism and growth. Besides colour change in the berries due to anthocyanin production and accumulation, berries soften as pectin and cellulose are degraded, acidity declines, and sugar accumulates. All these factors create a more favourable environment for microbial colonisation (Coombe, 1992; Renouf et al., 2005). Our study shows significant changes in the fungal diversity, and most distinct community structures of both the core (Fig. 3) and the whole (Supplementary Table S2) datasets at veraison. In particular, this observation is clearer in root zone soil and grape microbiome, when many non-core OTUs were only detected before or after veraison (Supplementary Table S3). Studies in other plants have also observed a shift in microbial communities during development, for example, Mougel et al. (2006) recorded that for *Medicago truncatula* the rhizosphere microbial communities differed according to the plant development stage, with the most significant shifts observed during the transition from vegetative to reproductive stages and to be strongly symbiotically associated. In viticulture, the microbial dynamics at veraison should be investigated to determine if there is a role for plant physiology, grape quality and resulting wine characteristics.

### The grapevine core microbiome responds to the changing environment

Over the annual growth cycle, grapevines are exposed to microorganisms from the surrounding environment (Stefanini and Cavalieri, 2018; Liu et al., 2019), and the microbiota are structured by the core microbiome (Fig. 4). Alongside the influences of changing metabolism during plant development, the core microbiome is also subject to dynamic environmental conditions where plants grow. Soil core microbiome closely correlated with the abiotic matrices (weather parameters; Supplementary Fig. S1), and it is in turn the biotic factor for the plant microbiome. Different plant organs respond to changes in their environment in different manners, via different and/or habitat-specific core taxa closely correlated with different environmental matrices (Fig. 5). Some important features distinguishing stages of soil microbiome identified by random forest models are also highly connected with plant core microbiome, including *Mortierella* sp., *Penicillium* sp., and *Ilyonectria* sp.. Water status (evaporation and relative soil moisture) and solar radiation had a strong influence on the microbiome. Higher solar radiation and temperature, decreased precipitation, increased evaporation and transpiration, and lower relative soil moisture were observed in the period from fruit set to veraison in both vineyards (data not shown). This water stress event coincided with dramatic changes in the microbiome discussed above. In viticulture, moderate water stress during ripening is recognised as beneficial for grape quality, resulting in lower yields and a concentration of berry metabolites (Van Leeuwen and Seguin, 2006). Our work may provide a microbial insight to understand additional mechanisms of how water status can affect quality wine production.

Other environmental factors, especially climate, structure microbial composition and biogeography across various habitats in soil and plant ecosystems (Fierer and Jackson, 2006; Martiny et al., 2006; Bokulich et al., 2014; Tedersoo et al., 2014; Burns et al., 2015). Our previous study revealed that wine-related fungal communities structured and distinguished vineyard ecosystems by impacting the flavour and quality of wine, and weather (in particular solar radiation and temperature) strongly affected soil and must fungal communities and the resultant wines across six winegrowing regions in southern Australia (Liu et al, under review). Vine water availability can condition the grape and must microbiota at regional scales, for example, precipitation and humidity correlates with the presence of filamentous fungi (for example *Penicillium*) (Bokulich et al., 2014) and yeasts (such as *Saccharomyces, Hanseniaspora*, and *Metschnikowia*) (Jara et al., 2016). Global studies show that climate (especially precipitation) (Tedersoo et al., 2014; Cavicchioli et al., 2019), followed by edaphic factors (such as soil pH, aridity) (Maestre et al., 2015; Bahram et al., 2018), drive soil fungal community assembly and hence multifunctionality in ecosystems (Bahram et al., 2018; Cavicchioli et al., 2019; Li et al., 2019). In the present study, microbial heterogeneity and seasonal dynamics across habitats highlight that water availability can influence the microbiota present within vineyards. Warming of vineyards and decline in soil water are the principal contributors to the effect of climate change on affecting grape growing and wine production (Webb et al., 2012). Grapevine phenology is shifting with climate change (Webb et al., 2012; Cameron et al., 2020), and our work suggests that the grapevine-associated microbiota will change as well. As the fungal microbiome associated with vineyards is a strong contributor to wine aroma and flavour, these changes will also affect the distinctiveness of regional wines (Liu et al., under review).

Microbial assemblages responding to climate change may also affect plant physiology, as reported for rhizosphere microorganisms which influence plant phenology and alter the flowering time (Wagner et al., 2014; Panke-Buisse et al., 2015; Lu et al., 2018). Interactions between grapevines and associated fungi deserve future study, and would provide further perspectives on vineyard establishment and management practices such as managing soil moisture and crop yield (Webb et al., 2012; Palliotti et al., 2014). As agricultural practices adapt to a changing climate, understanding the core microbiome, transient taxa and specific interactions between plant and microbe become all the more important to maintaining plant productivity (Cavicchioli et al., 2019).

## Conclusions

Our study describes the core microbiome of grapevines and illustrates the seasonal community succession driven by the core microbiome. Plants recruit their microorganisms based on habitats at different stages of development yet conserve core fungal species. While undertaken at two sites in one region in Australia, our research highlights the need to further investigate core microbial members for functions and interactions with the grapevine to understand the interactive functionalities that occur between the host crop and their microbiome. Increasing our understanding of fungal ecology in the vineyard and their responses to the changing environment over the course of the growing season would help improve management techniques to maintain healthy vines, produce quality grapes and allow expression of wine regionality.

## Experimental procedures

### Sample sites

Grapevine samples were collected at two vineyards in Mornington Peninsula wine region during the 2018 vintage (Fig. S2). KBS vineyard (KBS, 1.4 ha, altitude 55 m) and Merron’s vineyard (1.7 ha, altitude 195 m) are 5 km apart, planted with *Vitis vinifera* cv. Pinot Noir (clone MV6). Both vineyards are commercially operated by Stonier Wines under very similar viticultural practices, for example, grapevines were under vertical shoot positioning trellising systems and the same sprays were applied at the time of year. In each vineyard, five replicate vines were selected from the top, middle, and bottom of the dominant slope covering topological profiles of the vineyard, with pairwise distances ranging from 15 m to 135 m (Fig. S2A). GPS coordinates were recorded for each site and utilised to extract weekly weather data provided by Australian Water Availability Project (AWAP) (Jones et al., 2009) from two weather stations. Variables were observed by robust topography resolving analysis methods at a resolution of 0.05°× 0.05°(approximately 5 km × 5 km). Weekly measurements were extracted for precipitation (m), solar radiation (MJ/m^2^), total evaporation (m) (soil + vegetation), transpiration (m), mean high temperature (°C), mean low temperature (°C), mean temperature (°C), and mean relative soil moisture for each development stage during the growing season (October 2017 – April 2018).

### Sample collection

In each grapevine, four different grapevine habitats were studied: root zone soil, roots, leaves, and grapes or flowers (season dependent) (Fig. S2B). Samples were aseptically collected from the same vines at four developmental stages [as defined by Coombe (1992)] at flowering (November 2017), fruit-set (young berries enlarging, December 2017), veraison (berry colour change, January – February 2018), and harvest (berries ripe, March 2018) from both vineyards. Root zone soil was taken from the surrounding roots at the depth of 0 – 15 cm. Three sub-samples (the sun-exposed side, the shade side, and the middle) were mixed to form a composite sample. Healthy leaves of similar sizes were collected. Undamaged grapes were chosen from different grape bunches. Samples were immediately stored in sterile bags and transported to the laboratory on ice. Roots were collected and rinsed and cleaned of all visible soil using distilled water and flame-sterilized utensils back in the laboratory. Plant materials were flash frozen in liquid nitrogen and all the samples were stored at −80 °C until processing. A total of 160 vineyard samples were collected and processed.

### DNA extraction and sequencing

Genomic DNA was extracted from soil and botanical samples using PowerSoil™ DNA Isolation kits (QIAgen, CA, USA). DNA isolation of soil (0.25 g) samples followed the kit protocol. For plant parts, roots, leaves, grapes (removed seeds and stems) or flowers, were ground into powder in liquid nitrogen with 1% polyvinylpolypyrrolidone and then DNA isolated following the kit protocol. Thus, endophytes and epiphytes were extracted, sequenced, and analysed simultaneously from all plant organs. DNA extracts were stored at – 20 °C until further analysis.

Genomic DNA was submitted to Australian Genome Research Facility (AGRF) for amplification and sequencing. To assess the fungal communities, partial fungal internal transcribed spacer (ITS) region was amplified using the universal primer pairs ITS1F/2 (Gardes and Bruns, 1993). The primary PCR reactions contained 10 ng DNA template, 2× AmpliTaq Gold® 360 Master Mix (Life Technologies, Australia), 5 pmol of each primer. A secondary PCR to index the amplicons was performed with TaKaRa Taq DNA Polymerase (Clontech). Amplification was conducted under the following conditions: 95 °C for 7 min, followed by 35 cycles of 94 °C for 30 s, 55 °C for 45 s, 72 °C for 60 s and a final extension at 72 °C for 7 min. The resulting amplicons were cleaned again using magnetic beads, quantified by fluorometry (Promega Quantifluor) and normalised. The equimolar pool was cleaned a final time using magnetic beads to concentrate the pool and then measured using a High-Sensitivity D1000 Tape on an Agilent 2200 TapeStation. The pool was diluted to 5nM and molarity was confirmed again using a High-Sensitivity D1000 Tape.This was followed by 300 bp paired-end sequencing on an Illumina MiSeq (San Diego, CA, USA).

Raw sequences were processed using QIIME v1.9.2 (Caporaso et al., 2010). Low quality regions (Q < 20) were trimmed from the 5′ end of the sequences, and the paired ends were joined using FLASH (Magoc and Salzberg, 2011). Primers were trimmed and a further round of quality control was conducted to discard full length duplicate sequences, short sequences (< 100 nt), and sequences with ambiguous bases. Sequences were clustered followed by chimera checking using UCHIME algorithm from USEARCH v7.1.1090 (Edgar et al., 2011). Operational taxonomic units (OTUs) were assigned using UCLUST open-reference OTU-picking workflow with a threshold of 97% pairwise identity (Edgar, 2010). Singletons or unique reads in the resultant data set were discarded. Taxonomy was assigned to OTUs in QIIME using the UNITE fungal ITS database (v7.2) (Kõljalg et al., 2005). To avoid/ reduce biases generated by varying sequencing depth, sequences were rarefied to 10,000 per sample (the lowest sequencing depth) prior to downstream analysis. Raw sequencing reads have been deposited at the National Centre for Biotechnology Information Sequence Read Archive under the bioproject PRJNA626608.

### Data analysis

Fungal OTUs shared among habitats in the vineyard was illustrated by the “VennDiagram” package (Chen and Boutros, 2011) in R (v3.5.0). Microbial alpha-diversity was calculated using the Shannon index with the “vegan” package (Oksanen et al., 2007). One-way analysis of variance (ANOVA) was used to determine whether sample classifications (e.g., habitat, developmental stage, vineyard) contained statistically significant differences in the diversity. Principal coordinate analysis (PCoA) was performed to evaluate the distribution patterns of grapevine-associated microbiome based on beta-diversity calculated by the Bray–Curtis distance with the “labdsv” package (Roberts, 2007). Permutational multivariate analysis of variance using distance matrices with 999 permutations was conducted within each sample category to determine the statistically significant differences with “adonis” function in “vegan” (Oksanen et al., 2007). Significant taxonomic differences of fungi between vineyard habitats were tested using linear discriminant analysis (LDA) effect size (LEfSe) analysis (Segata et al., 2011) (https://huttenhower.sph.harvard.edu/galaxy/). The OTU table was filtered to include only OTUs > 0.1% relative abundance to reduce LEfSe complexity. The factorial Kruskal– Wallis sum-rank test (*α* = 0.05) was applied to identify taxa with significant differential abundances between classes (all-against-all comparisons), followed by the logarithmic LDA score (threshold = 2.0) to estimate the effect size of each discriminative feature. Significant taxa were used to generate taxonomic cladograms illustrating differences between sample categories.

The core microbiome was defined based on abundance-occupancy distributions that include highly abundant and ubiquitous taxa (Delgado-Baquerizo et al., 2018; Shade and Stopnisek, 2019). For soil core microbiome, OTUs were filtered out with top 1% mean relative abundance (MRA) across all samples (Galand et al., 2009) and an occurrence frequency found in more than 50% of samples or 100% occupancy at any time point (Delgado-Baquerizo et al., 2018; Grady et al., 2019). For the grapevine, OTUs with top 10% MRA across all samples (roots, leaves, grapes, flowers) and 50% occupancy of all samples or 100% occupancy of one habitat samples were defined as the core members (Delgado-Baquerizo et al., 2018; Grady et al., 2019). For each plant organ (except flowers), the core OTUs were those top 10% abundant across all samples with 50% occupancy of all samples or 100% occupancy at any developmental stage (Delgado-Baquerizo et al., 2018; Grady et al., 2019). Dynamics and successions of the core microbiome were illustrated by alluvial diagrams using the “ggalluvial” package (Brunson, 2018) in “ggplot2”.

Random forest supervised classification models (Breiman, 2001) were used to evaluate the discriminative power of fungal microbiota (all OTUs) to distinguish developmental stages and the robustness of the groupings themselves. The importance of each predictor is measured by the mean decrease in Gini coefficient (MDG) (that is, how each variable contributes to the homogeneity of the nodes and leaves in the resulting random forest, and variables resulting in nodes with higher purity have a higher MDG). Models were constructed with 10,000 trees, with OTUs as predictors and developmental stages as class labels using the “randomForest” package (Liaw and Wiener, 2002).

The co-occurrence/interaction patterns between the core microbiome associated with grapevine habitats (grapes, leaves, roots) and soil, and weather conditions during the growing season was explored using network analysis with Cytoscape (Shannon et al., 2003). Correlation matrices were calculated with all possible pair-wise Spearman’s rank correlations between selected OTUs of plant and soil, or OTUs and weather indexes. Correlations with a Spearman correlation coefficient *ρ* ≥ 0.8 and a *p* < 0.01 were considered statistically robust and displayed in the networks (Junker and Schreiber, 2008).

## Acknowledgements

We give our sincere thanks to Stonier Wines in Mornington Peninsula who kindly allowed vineyard access and enabled sampling. DL acknowledges support from a Ph.D. scholarship and funding from Wine Australia (AGW Ph1602) and a Melbourne Research Scholarship from the University of Melbourne. Dr Qinling Chen, Prof Deli Chen and Dr Pangzhen Zhang are acknowledged for helpful discussions and support in the course of this work.

## Supplementary tables and figures

**Table S1.**
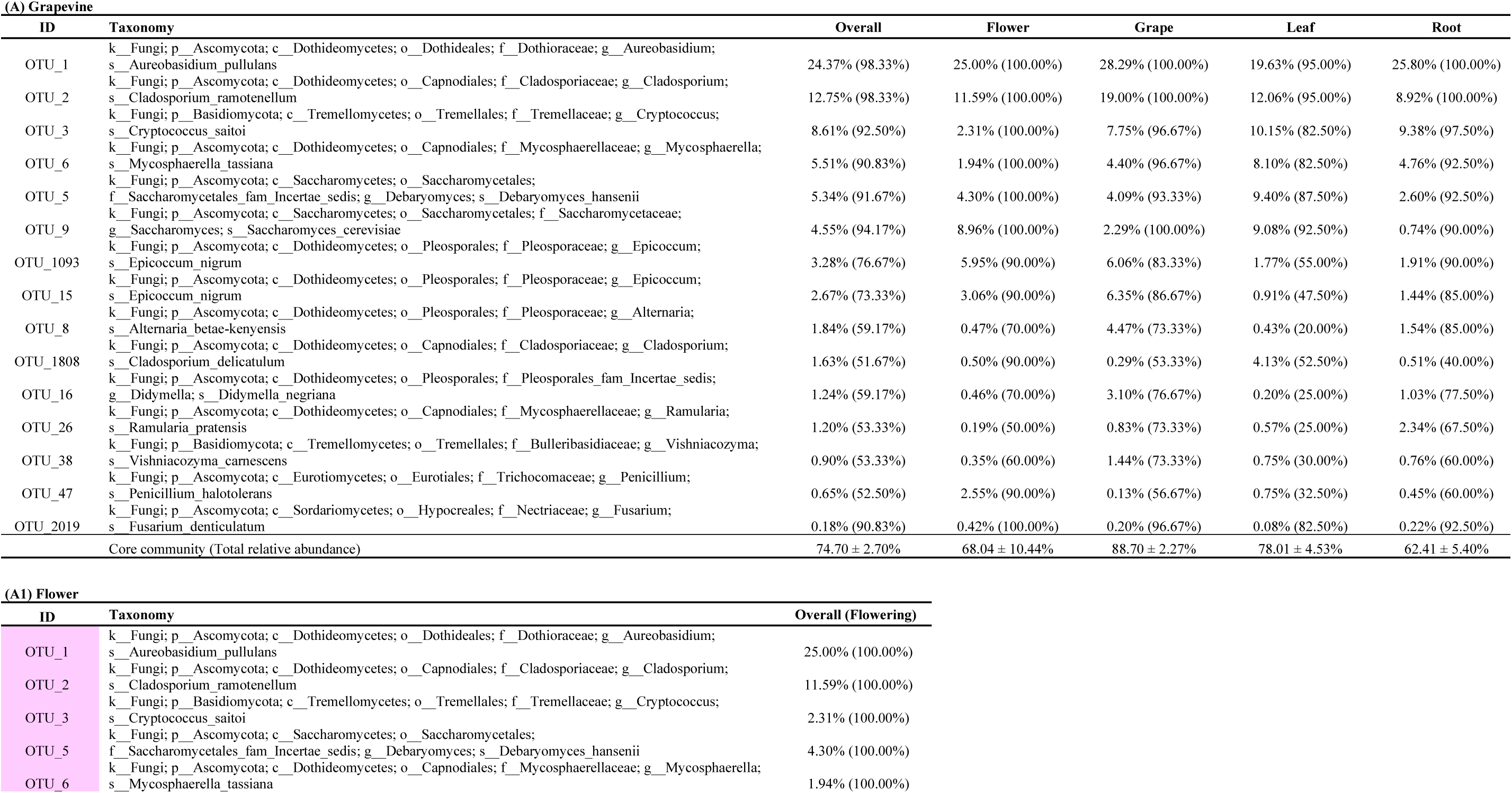
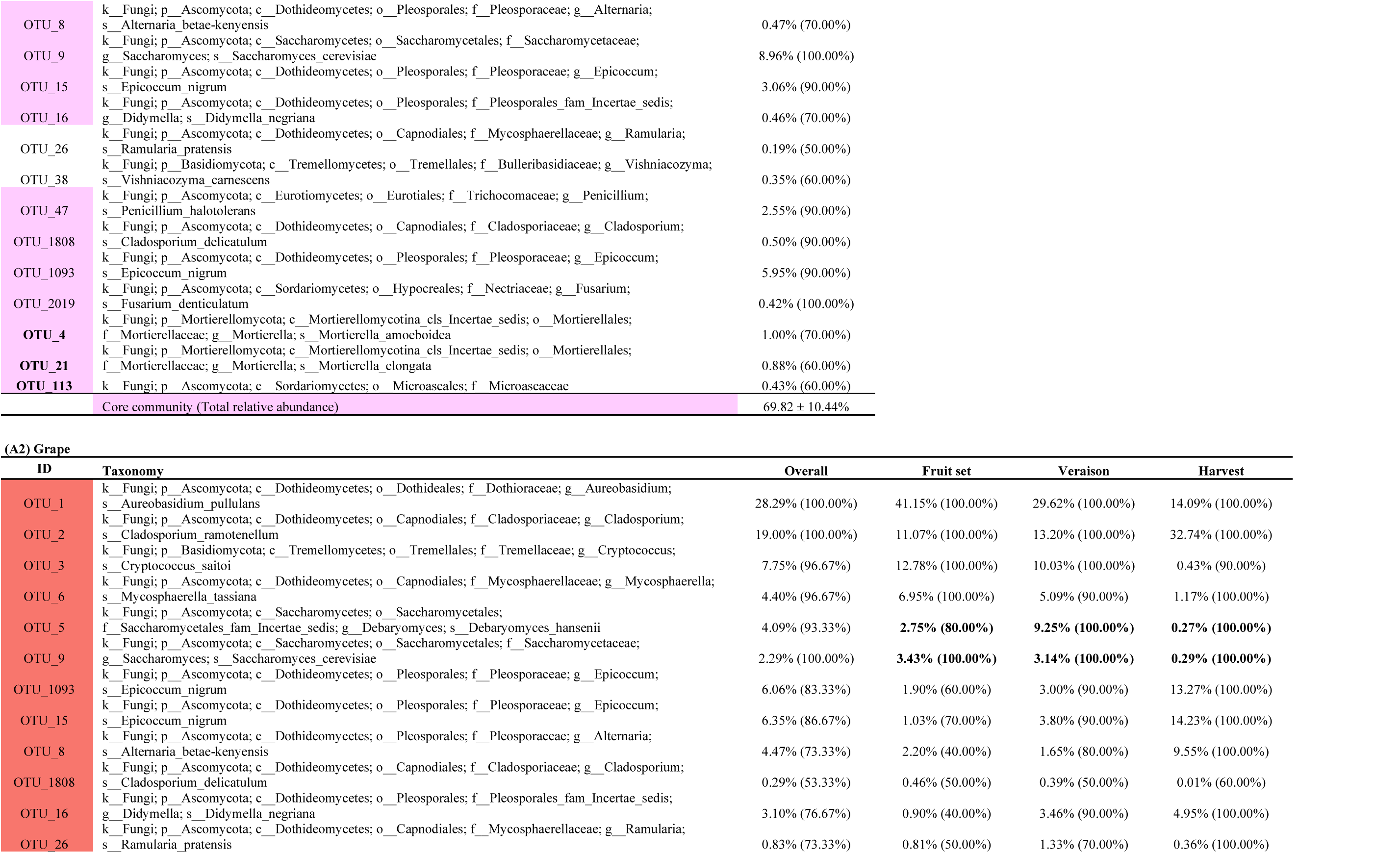
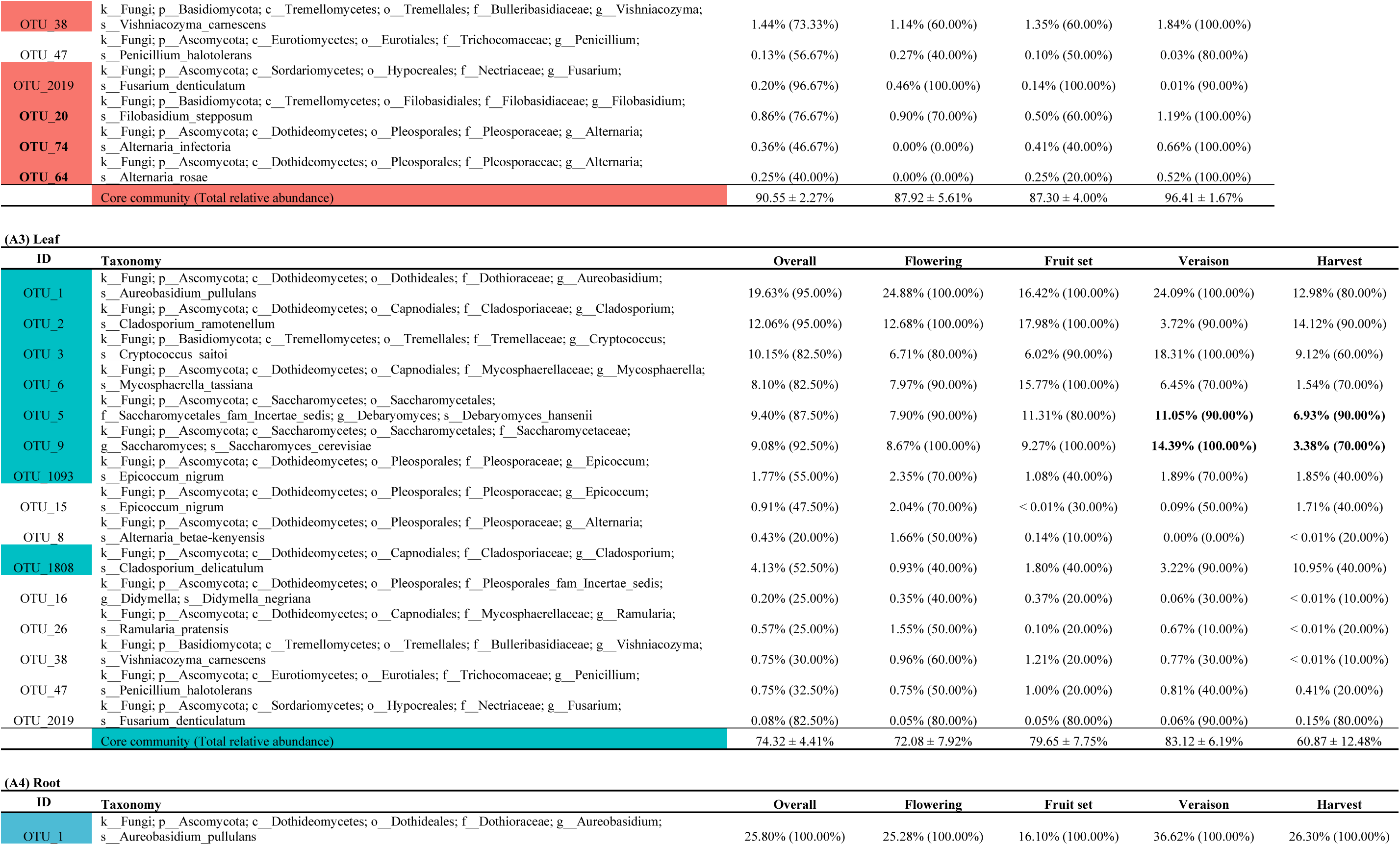
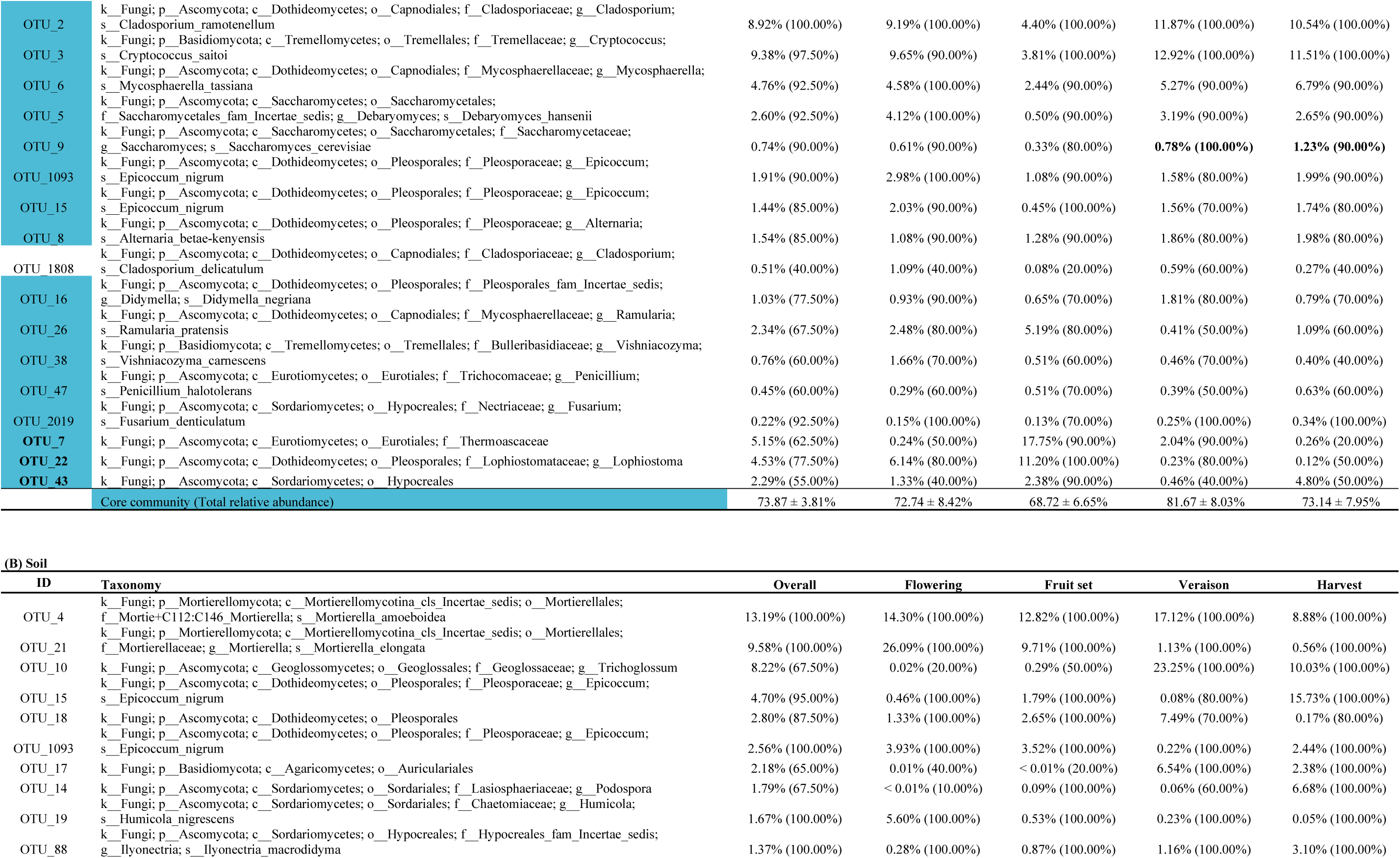
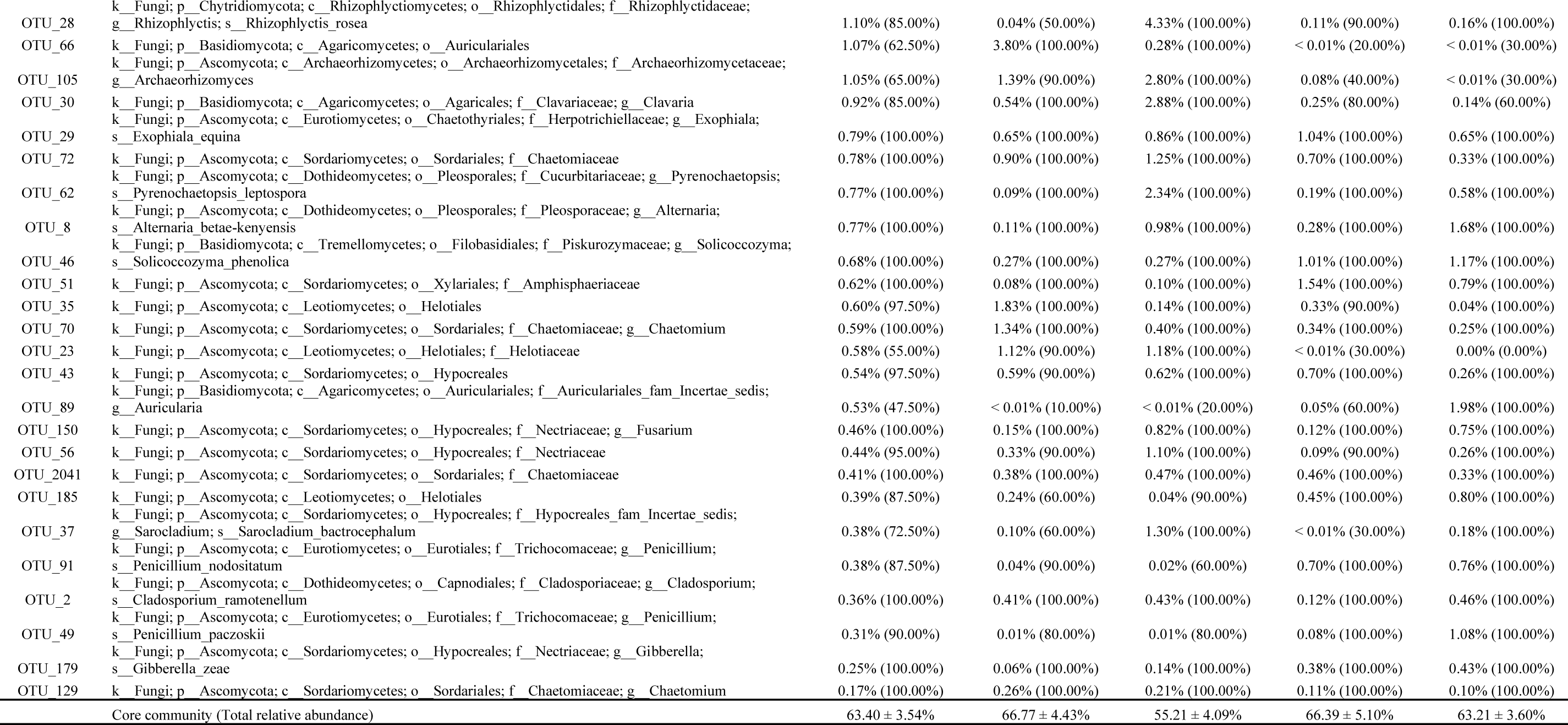
OTUs identified as core community assoicated with the grapevine or within each vineyard habitat The first column shows the OTU id and the second column shows the lowest taxonomy level the OTU could be classified to. The third to the last column show the mean relative abundance (outside the parentheses) and occupancy of samples (inside the parentheses) across plant organs(A) or within each vineyard habitat(A1 – A4, grapevine; B soil). Core OTUs are coloured for each plant organ, of which bold ones are core OTUs that are specific to this organ. For example, core OTUs of grapes are those filled with red, of which OTU_20, OTU_74, OTU_64 are grape-specific core OTUs.

**Table S2.** Classification of random forest models of the development stage

|  |  | Flowering | Fruit set | Veraison | Harvest | Class Error |
| --- | --- | --- | --- | --- | --- | --- |
| Grape | Fruit set | N.A. | 9 | 1 | 0 | 0.100 |
|  | Veraison | N.A. | 1 | 9 | 0 | 0.100 |
|  | Harvest | N.A. | 0 | 1 | 9 | 0.100 |
| Leaf | Flowering | 6 | 2 | 1 | 1 | 0.400 |
|  | Fruit set | 1 | 6 | 0 | 3 | 0.400 |
|  | Veraison | 0 | 2 | 7 | 1 | 0.300 |
|  | Harvest | 0 | 1 | 3 | 6 | 0.400 |
| Root | Flowering | 8 | 2 | 0 | 0 | 0.200 |
|  | Fruit set | 3 | 7 | 0 | 0 | 0.400 |
|  | Veraison | 0 | 0 | 9 | 1 | 0.200 |
|  | Harvest | 0 | 0 | 3 | 7 | 0.300 |
| Soil | Flowering | 9 | 1 | 0 | 0 | 0.100 |
|  | Fruit set | 3 | 7 | 0 | 0 | 0.300 |
|  | Veraison | 0 | 0 | 10 | 0 | 0.000 |
|  | Harvest | 0 | 0 | 1 | 9 | 0.100 |

N.A.: Not applied

**Table S3.** Important features of random forest models outside the core community The same as Table S1.

| (A) Grape |  |  |  |  |  |  |
| --- | --- | --- | --- | --- | --- | --- |
| ID | Taxonomy | Overall | Fruit set | Veraison | Harvest |  |
| OTU_146 | k__Fungi; p__Ascomycota; c__Dothideomycetes; o__Capnodiales; f__Capnodiales_fam_Incertae_sedis; g__Rachicladosporium; s__Rachicladosporium_cboliae | 0.04% (30.00%) | 0.00% (0.00%) | 0.00% (0.00%) | 0.12% (90.00%) |  |
| OTU_54 | k__Fungi; p__Ascomycota; c__Dothideomycetes; o__Pleosporales; f__Pleosporaceae; g__Stemphylium; s__Stemphylium_herbarum | 0.51% (46.67%) | 0.00% (0.00%) | 0.70% (40.00%) | 0.83% (100.00%) |  |
| OTU_33 | k__Fungi; p__Ascomycota; c__Saccharomycetes; o__Saccharomycetales; f__Saccharomycodaceae; g__Hanseniaspora; s__Hanseniaspora_uvarum | 0.06% (36.67%) | 0.04% (20.00%) | 0.11% (50.00%) | 0.02% (40.00%) |  |
| OTU_40 | k__Fungi; p__Basidiomycota; c__Microbotryomycetes; o__Sporidiobolales; f__Sporidiobolales_fam_Incertae_sedis; g__Rhodotorula; s__Rhodotorula_babjevae | 0.44% (30.00%) | 0.67% (10.00%) | 0.23% (10.00%) | 0.41% (70.00%) |  |
| OTU_37 | k__Fungi; p__Ascomycota; c__Sordariomycetes; o__Hypocreales; f__Hypocreales_fam_Incertae_sedis; g__Sarocladium; s__Sarocladium_bactrocephalum | 0.06% (23.33%) | 0.12% (10.00%) | 0.00% (0.00%) | 0.06% (60.00%) |  |
| OTU_2201 | k__Fungi; p__Ascomycota; c__Sordariomycetes; o__Hypocreales; f__Nectriaceae; g__Fusarium; s__Fusarium_denticulatum | 0.05% (46.67%) | 0.01% (70.00%) | 0.12% (70.00%) | 0.00% (0.00%) |  |
| OTU_121 | k__Fungi; p__Ascomycota; c__Saccharomycetes; o__Saccharomycetales; f__Saccharomycetaceae; g__Torulaspora; s__Torulaspora_delbrueckii | 0.08% (36.67%) | 0.00% (0.00%) | 0.23% (50.00%) | 0.01% (60.00%) |  |
| OTU_4 | k__Fungi; p__Mortierellomycota; c__Mortierellomycotina_cls_Incertae_sedis; o__Mortierellales; f__Mortierellaceae; g__Mortierella; s__Mortierella_amoeboidea | 0.52% (26.67%) | 0.83% (30.00%) | 0.72% (40.00%) | < 0.01% (10.00%) |  |
| OTU_323 | k__Fungi; p__Ascomycota; c__Dothideomycetes; o__Pleosporales; f__Pleosporales_fam_Incertae_sedis; g__Didymella; s__Didymella_negriana | 0.20% (30.00%) | 0.00% (0.00%) | 0.24% (50.00%) | 0.36% (40.00%) |  |
| OTU_128 | k__Fungi; p__Ascomycota; c__Dothideomycetes; o__Capnodiales; f__Cladosporiaceae; g__Cladosporium; s__Cladosporium_sphaerospermum | 0.11% (30.00%) | 0.00% (0.00%) | 0.33% (40.00%) | 0.01% (50.00%) |  |
| (B) Leaf |  |  |  |  |  |  |
| ID | Taxonomy | Overall | Flowering | Fruit set | Veraison | Harvest |
| OTU_21 | k__Fungi; p__Mortierellomycota; c__Mortierellomycotina_cls_Incertae_sedis; o__Mortierellales; f__Mortierellaceae; g__Mortierella; s__Mortierella_elongata | 0.04% (20.00%) | 0.15% (10.00%) | < 0.01% (10.00%) | < 0.01% (10.00%) | 0.01% (50.00%) |
| OTU_48 | k__Fungi; p__Ascomycota; c__Dothideomycetes; o__Pleosporales; f__Pleosporales_fam_Incertae_sedis; g__Boeremia; s__Boeremia_noackiana | 2.74% (17.50%) | 0.14% (20.00%) | 0.00% (0.00%) | < 0.01% (10.00%) | 11.10% (40.00%) |
| OTU_4 | k__Fungi; p__Mortierellomycota; c__Mortierellomycotina_cls_Incertae_sedis; o__Mortierellales; f__Mortierellaceae; g__Mortierella; s__Mortierella_amoeboidea | 0.26% (27.50%) | 0.91% (30.00%) | 0.00% (0.00%) | 0.12% (30.00%) | 0.02% (50.00%) |
| OTU_2078 | k__Fungi; p__Ascomycota; c__Dothideomycetes; o__Capnodiales; f__Cladosporiaceae; g__Cladosporium; s__Cladosporium_delicatulum | 0.01% (30.00%) | < 0.01% (20.00%) | < 0.01% (30.00%) | < 0.01% (50.00%) | 0.01% (20.00%) |
| OTU_24 | k__Fungi; p__Ascomycota; c__Eurotiomycetes; o__Eurotiales; f__Trichocomaceae; g__Aspergillus; s__Aspergillus_conicus | 1.96% (15.00%) | 0.11% (30.00%) | 6.77% (10.00%) | 0.38% (20.00%) | 0.00% (0.00%) |
| OTU_20 | k__Fungi; p__Basidiomycota; c__Tremellomycetes; o__Filobasidiales; f__Filobasidiaceae; g__Filobasidium; s__Filobasidium_stepposum | 1.42% (25.00%) | 1.64% (30.00%) | 0.25% (20.00%) | 1.19% (40.00%) | 2.58% (10.00%) |
| OTU_2201 | k__Fungi; p__Ascomycota; c__Sordariomycetes; o__Hypocreales; f__Nectriaceae; g__Fusarium; s__Fusarium_denticulatum | < 0.01% (35.00%) | < 0.01% (20.00%) | < 0.01% (30.00%) | < 0.01% (40.00%) | < 0.01% (50.00%) |
| OTU_23 | k__Fungi; p__Ascomycota; c__Leotiomycetes; o__Helotiales; f__Helotiaceae | < 0.01% (10.00%) | 0.00% (0.00%) | 0.00% (0.00%) | 0.00% (0.00%) | 0.01% (40.00%) |
| OTU_323 | k__Fungi; p__Ascomycota; c__Dothideomycetes; o__Pleosporales; f__Pleosporales_fam_Incertae_sedis; g__Didymella; s__Didymella_negriana | < 0.01% (5.00%) | 0.00% (0.00%) | < 0.01% (20.00%) | 0.00% (0.00%) | 0.00% (0.00%) |
| OTU_35 | k__Fungi; p__Ascomycota; c__Leotiomycetes; o__Helotiales | 0.65% (5.00%) | 2.68% (20.00%) | 0.00% (0.00%) | 0.00% (0.00%) | 0.00% (0.00%) |
| OTU_121 | k__Fungi; p__Ascomycota; c__Saccharomycetes; o__Saccharomycetales; f__Saccharomycetaceae; g__Torulaspora; s__Torulaspora_delbrueckii | 0.21% (17.50%) | 0.78% (20.00%) | 0.02% (30.00%) | 0.07% (20.00%) | 0.00% (0.00%) |
| OTU_33 | k__Fungi; p__Ascomycota; c__Saccharomycetes; o__Saccharomycetales; f__Saccharomycodaceae; g__Hanseniaspora; s__Hanseniaspora_uvarum | 2.15% (25.00%) | 0.69% (20.00%) | 1.25% (30.00%) | 1.33% (20.00%) | 5.29% (30.00%) |

| ID | Taxonomy | Overall | Flowering | Fruit set | Veraison | Harvest |
| --- | --- | --- | --- | --- | --- | --- |
| OTU_17 | k__Fungi; p__Basidiomycota; c__Agaricomycetes; o__Auriculariales | 0.80% (22.50%) | 2.70% (40.00%) | 0.41% (50.00%) | 0.00% (0.00%) | 0.00% (0.00%) |
| OTU_4 | k__Fungi; p__Mortierellomycota; c__Mortierellomycotina_cls_Incertae_sedis; o__Mortierellales; f__Mortierellaceae; g__Mortierella; s__Mortierella_amoeboida | 0.67% (45.00%) | 0.00% (0.00%) | 0.55% (90.00%) | 0.91% (40.00%) | 1.25% (50.00%) |
| OTU_51 | k__Fungi; p__Ascomycota; c__Sordariomycetes; o__Xylariales; f__Amphisphaeriaceae | 0.09% (22.50%) | 0.00% (0.00%) | 0.08% (40.00%) | 0.00% (0.00%) | 0.29% (50.00%) |
| OTU_122 | k__Fungi; p__Ascomycota; c__Sordariomycetes; o__Hypocreales; f__Bionectriaceae; g__Clonostachys; s__Clonostachys_rosea | 0.34% (32.50%) | 0.04% (30.00%) | 0.27% (50.00%) | 0.00% (0.00%) | 1.01% (50.00%) |
| OTU_33 | k__Fungi; p__Ascomycota; c__Saccharomycetes; o__Saccharomycetales; f__Saccharomycodaceae; g__Hanseniaspora; s__Hanseniaspora_uvarum | 0.03% (40.00%) | 0.02% (40.00%) | 0.02% (50.00%) | < 0.01% (30.00%) | 0.08% (40.00%) |
| OTU_29 | k__Fungi; p__Ascomycota; c__Eurotiomycetes; o__Chaetothyriales; f__Herpotrichiellaceae; g__Exophiala; s__Exophiala_equina | 0.25% (32.50%) | 0.30% (20.00%) | 0.42% (80.00%) | 0.00% (0.00%) | 0.23% (30.00%) |
| OTU_20 | k__Fungi; p__Basidiomycota; c__Tremellomycetes; o__Filobasidiales; f__Filobasidiaceae; g__Filobasidium; s__Filobasidium_stepposum | 0.18% (20.00%) | 0.57% (50.00%) | < 0.01% (10.00%) | 0.00% (0.00%) | 0.14% (20.00%) |
| OTU_2201 | k__Fungi; p__Ascomycota; c__Sordariomycetes; o__Hypocreales; f__Nectriaceae; g__Fusarium; s__Fusarium_denticulatum | 0.01% (52.50%) | 0.01% (60.00%) | < 0.01% (40.00%) | < 0.01% (30.00%) | 0.01% (80.00%) |

(D) Soil
| ID | Taxonomy | Overall | Flowering | Fruit set | Veraison | Harvest |
| --- | --- | --- | --- | --- | --- | --- |
| OTU_584 | k__Fungi; p__Ascomycota; c__Eurotiomycetes; o__Eurotiales; f__Trichocomaceae; g__Penicillium; s__Penicillium_multicolor | 0.01% (32.50%) | 0.00% (0.00%) | 0.00% (0.00%) | < 0.01% (20.00%) | 0.02% (100.00%) |
| OTU_501 | k__Fungi; p__Ascomycota; c__Dothideomycetes; o__Pleosporales; f__Pleosporales_fam_Incertae_sedis; g__Didymella; s__Didymella_pedeiae | 0.14% (75.00%) | 0.03% (30.00%) | 0.03% (70.00%) | 0.12% (100.00%) | 0.36% (100.00%) |
| OTU_151 | k__Fungi; p__Ascomycota; c__Leotiomycetes; o__Helotiales | 0.11% (42.50%) | 0.42% (100.00%) | < 0.01% (60.00%) | 0.00% (0.00%) | 0.00% (0.00%) |
| OTU_519 | k__Fungi; p__Ascomycota; c__Archaeorhizomycetes; o__Archaeorhizomycetales; f__Archaeorhizomycetaceae; g__Archaeorhizomyces | 0.03% (57.50%) | 0.06% (100.00%) | 0.07% (100.00%) | < 0.01% (20.00%) | 0.00% (0.00%) |
| OTU_763 | k__Fungi; p__Mortierellomycota; c__Mortierellomycotina_cls_Incertae_sedis; o__Mortierellales; f__Mortierellaceae; g__Mortierella; s__Mortierella_elongata | 0.13% (60.00%) | 0.41% (100.00%) | 0.09% (90.00%) | 0.01% (30.00%) | < 0.01% (10.00%) |
| OTU_223 | k__Fungi; p__Basidiomycota; c__Tremellomycetes; o__Filobasidiales; f__Piskurozymaceae; g__Solicoccozyma; s__Solicoccozyma_terricola | 0.04% (65.00%) | 0.07% (100.00%) | 0.09% (100.00%) | < 0.01% (20.00%) | < 0.01% (30.00%) |
| OTU_392 | k__Fungi; p__Ascomycota; c__Lecanoromycetes; o__Lecanorales; f__Cladoniaceae; g__Cladonia; s__Cladonia_cervicornis | 0.01% (45.00%) | 0.02% (70.00%) | 0.03% (100.00%) | 0.00% (0.00%) | 0.00% (0.00%) |
| OTU_94 | k__Fungi; p__Ascomycota; c__Eurotiomycetes; o__Eurotiales; f__Trichocomaceae; g__Aspergillus; s__Aspergillus_ardalensis | 0.01% (60.00%) | < 0.01% (20.00%) | < 0.01% (30.00%) | 0.02% (80.00%) | 0.03% (100.00%) |
| OTU_341 | k__Fungi; p__Ascomycota; c__Sordariomycetes; o__Sordariales; f__Lasiosphaeriaceae; g__Podospora | 0.05% (50.00%) | 0.00% (0.00%) | 0.00% (0.00%) | 0.08% (80.00%) | 0.12% (100.00%) |

**Table S4.** ANOVA results of weather conditions between developmental stages

| Factor | Weather parameter | Df | F | *p* | Pr (>F) |
| --- | --- | --- | --- | --- | --- |
| Env1 | Solar radiation | 3 | 98.42 | 3.33E-04 | *** |
| Env2 | Mean high temperature | 3 | 49.88 | 0.002 | ** |
| Env3 | Mean low temperature | 3 | 165.4 | 1.20E-04 | *** |
| Env4 | Mean temperature | 3 | 29.72 | 0.003 | ** |
| Env5 | Precipitation | 3 | 87.13 | 4.24E-04 | *** |
| Env6 | Evaporation | 3 | 279.6 | 4.22E-05 | *** |
| Env7 | Transpiration | 3 | 635.5 | 8.21E-06 | *** |
| Env8 | Relative soil moisture | 3 | 118.2 | 2.32E-04 | *** |

**Fig. S1.**
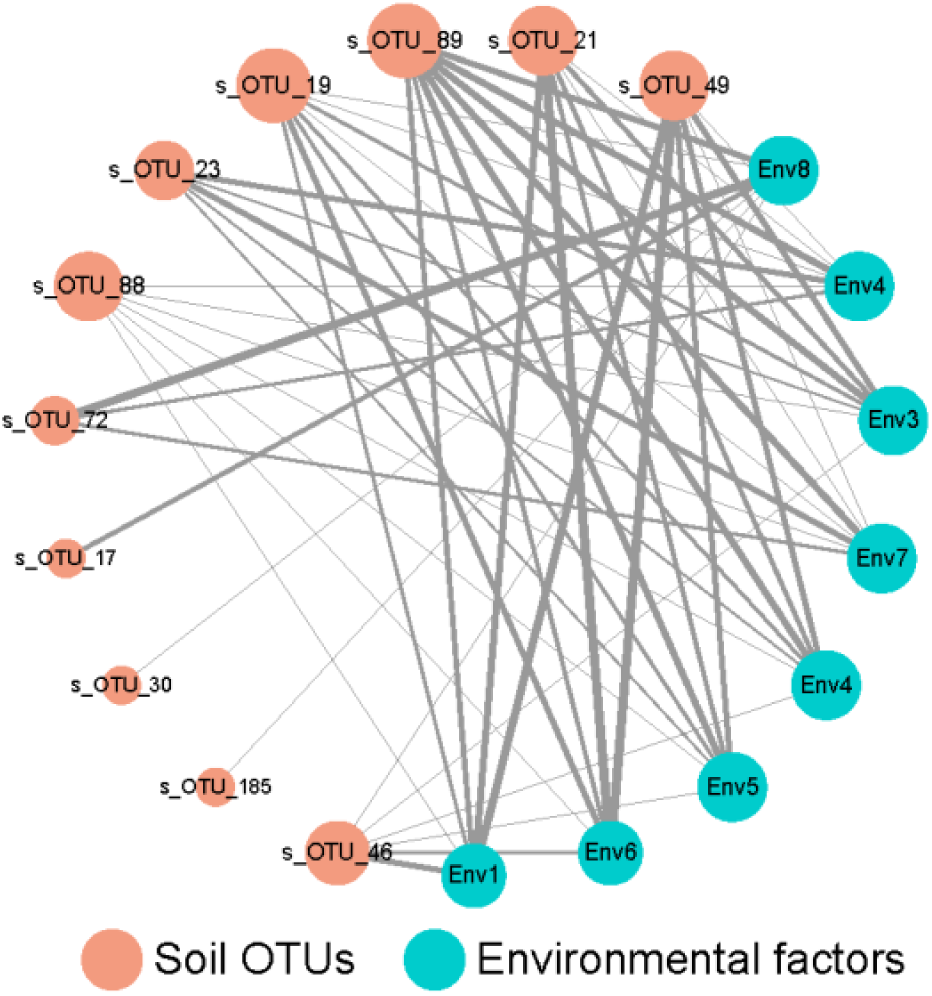
Co-occurrence pattern between the soil core microbiome and weather parameters. Circle nodes represent soil core OTUs and environmental variables, with different colours. Direct connections between nodes indicate strong correlations (Spearman correlation coefficient, ρ ≥ 0.8; p < 0.01). The size of nodes is proportioned to the interconnected degree. The width of edges is proportioned to the correlation coefficients. Abbreviations: Env1, solar radiation; Env2, mean high temperature; Env3, mean low temperature; Env4, mean temperature; Env5, precipitation; Env6, evaporation; Env7, transpiration; Env8, relative soil moisture.

**Fig. S2.**
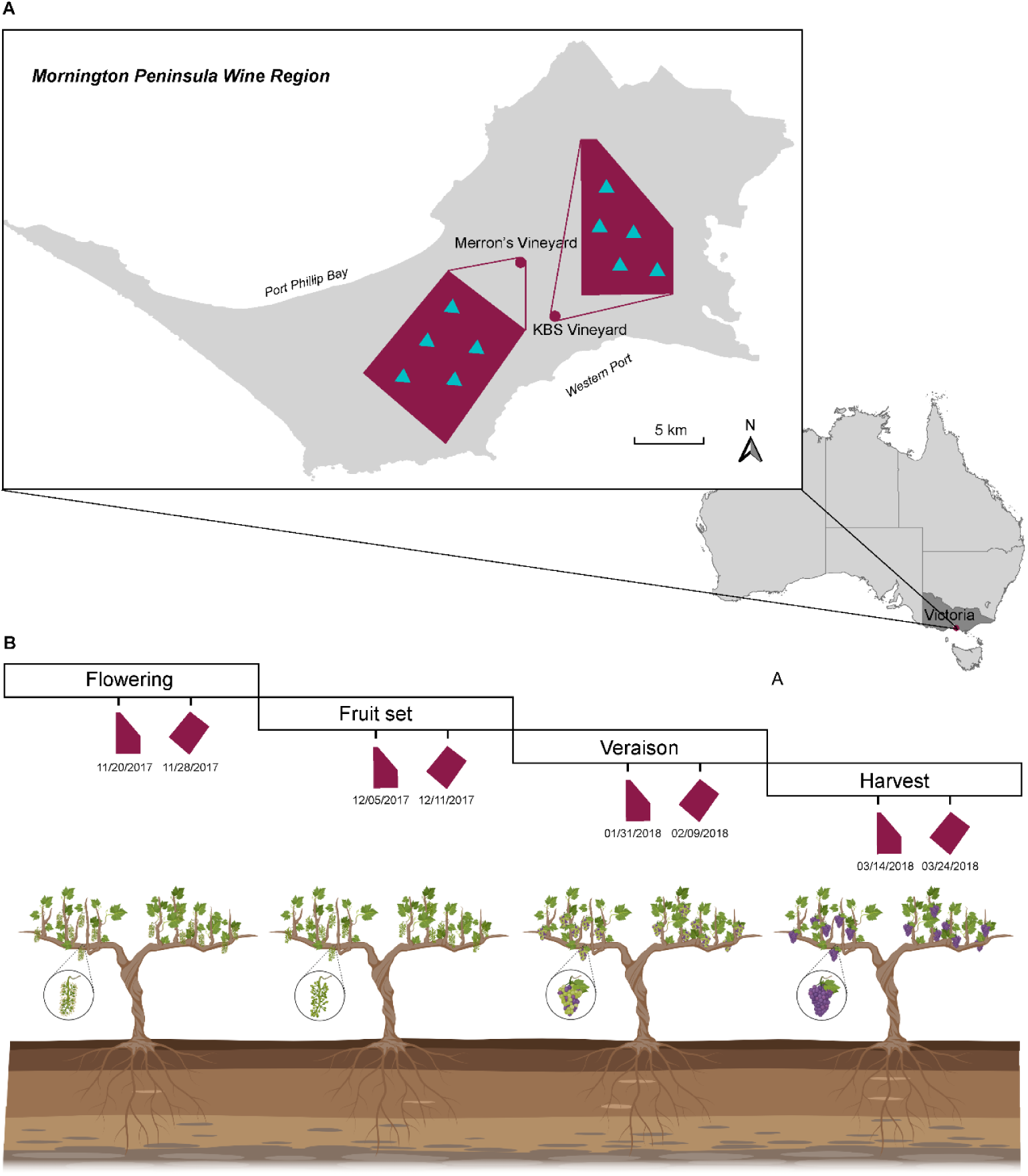
Experimental design and sampling scheme. (A) Sampling map of two vineyards in Mornington Peninsula wine region, Australia. Five *Vitis vinifera* Pinot Noir vines were selected in each vineyard. (B) Soils, roots, leaves, and grapes/ flowers were collected in each grapevine throughout the annual growth cycle: flowering, fruit set, veraison (berry colour change), and harvest. Calendar dates of sample collection are indicated for each developmental stage.

